# Sampling bias and the robustness of ecological metrics for plant-damage-type association networks

**DOI:** 10.1101/2022.07.23.501238

**Authors:** Anshuman Swain, Lauren E. Azevedo-Schmidt, S. Augusta Maccracken, Ellen D. Currano, Jennifer Dunne, Conrad C. Labandeira, William F. Fagan

## Abstract

Plants and their insect herbivores have been a dominant component of the terrestrial ecological landscape for the past 410 million years and feature intricate evolutionary patterns and co-dependencies. A complex systems perspective allows for both detailed resolution of these evolutionary relationships as well as comparison and synthesis across systems. Using proxy data of insect herbivore damage (denoted by the damage type or DT) preserved on fossil leaves, functional bipartite network representations provide insights into how plant–insect associations depend on geological time, paleogeographical space, and environmental variables such as temperature and precipitation. However, the metrics measured from such networks are prone to sampling bias. Such sensitivity is of special concern for plant–DT association networks in paleontological settings where sampling effort is often severely limited. Here, we explore the sensitivity of functional bipartite network metrics to sampling intensity and identify sampling thresholds above which metrics appear robust to sampling effort. Across a broad range of sampling efforts, we find network metrics to be less affected by sampling bias and/or sample size than richness metrics, which are routinely used in studies of fossil plant–DT interactions. These results provide reassurance that cross-comparisons of plant–DT networks offer insights into network structure and function and support their widespread use in paleoecology. Moreover, these findings suggest novel opportunities for using plant–DT networks in neontological terrestrial ecology to understand functional aspects of insect herbivory across geological time, environmental perturbations, and geographic space.

## Introduction

Ecological interactions among species come in many forms, are extremely complex, and are spatiotemporally variable. Researchers trying to reconstruct such interactions through limited observations often focus on a set of species of interest (e.g., Dormann et al., 2017; Agrawal et al., 2012; Dyer et al., 2007). To understand and explore their complex interdependencies at an ecosystem scale, these interactions often are represented as an ecological network involving two specific sets of biological entities, such as plant hosts and their dependent insect herbivore species and pathogens. Such two-part networks are called bipartite networks (Bascompte and Jordano, 2013; Dormann et al., 2017), and researchers often focus on taxonomic bipartite networks where nodes are Linnaean taxa. Taxonomic bipartite networks have proved instrumental in elucidating numerous important aspects of various ecological processes (Bascompte and Jordano, 2013) and are widely used for network studies in modern ecology.

A shift from taxonomic, or species-specific studies, to functional studies, especially studies involving functional traits, has shown the importance of ecological drivers on interactions in both the modern and paleontological ecological literature (e.g., Kunstler et al., 2016; Perez et al., 2020; Barnosky et al., 2017; Schellenberger Costa et al., 2018). Thus, the progression from taxonomic to functional bipartite network analyses can allow for the comparison of interacting trophic levels regardless of taxonomic classification, the designation of which is often challenging or inaccessible within paleontological, deep-time studies. Additionally, comparisons among modern taxonomic bipartite networks are often difficult due to the presence of disjoint species/taxonomic units for both classes of nodes, and most comparisons rely solely on overall network metrics.

In this work, we focus primarily on plant–insect herbivore associations. Identification of the insect herbivore species is essential for building taxonomic bipartite networks (which is the case for all modern herbivory networks), but such information is seldom available for the fossil record. Therefore, taxonomic networks are often impossible to construct in palaeoecological cases. Additionally, as mentioned above, comparisons between modern taxonomic networks of insect herbivory are difficult due to high insect (and plant) species turnover across space. Both of these problems can be tackled to a substantial extent by using a functional approach to insect herbivory rather than a taxonomic approach, and such an approach is not only useful for paleoecological insect herbivory studies but also for cross-comparisons among modern ones.

The node classes in functional bipartite networks that can be used for such studies are plant hosts (often determined using compression leaf fossils in palaeoecological cases), and insect/arthropod or pathogen damage (often termed as damage types). Insect feeding damage or damage types (DTs) are morphologically distinct, stereotyped, feeding patterns preserved on plant tissue, which have remained almost unchanged in structure over large periods of geological time and geographical space (Labandeira and Wappler, 2022), making them a good functional representative of insect herbivory and other damages on plants. DTs are the most abundant available data on ancient herbivorous insects and their ecological associations, whereas insect body fossils are sporadic through the geologic record (Labandeira & Eble 2000) and do not often preserve information regarding their ecological associations (i.e., plant–insect interactions). Direct observations of ancient insects feeding upon specific plants are rarer still, such as scale insects preserved on fossil leaves and stems (Xiao et al. 2021a). These functional bipartite networks are a paleoecological counterpart, or analog, to the taxonomically based plant–insect interaction networks commonly found in the modern ecological literature (Table 1; major differences between taxonomic versus functional bipartite networks). Examples from the modern literature include undirected network associations between woody perennial species and their pathogens, specialization in plant host–gall interactions, and the preference of insects for particular fern hosts (Fodor and Hâruţa, 2014; Araújo and Kollár, 2019; Fuentes-Jacques et al., 2020). However, for fossil data, one of these node classes, the DTs functionally serve as ecological units that have links to plant taxa and would incorporate data such as one herbivore species producing multiple DTs and multiple herbivore species producing one DT, the multidamagers and monodamagers of Carvalho et al. (2014). Such a formulation would allow comparisons of functional bipartite networks to be potentially useful in the study of deep time plant–insect associations, such as the evolution of insect feeding strategies across considerable time intervals, up to and including present day ecosystems.

**Table 1:** Comparison of taxonomic versus functional bipartite networks

| **<u>Feature</u>** | **<u>Taxonomic bipartite network</u>** | **<u>Functional bipartite network</u>** |
| --- | --- | --- |
| **Node classes** | Plant host taxa; herbivore taxa | Plant host taxa; plant–arthropod/pathogen interaction (damage types/DT) attributes |
| **Variables for node classes** | Taxa (e.g., species, genera) for both nodes | Taxa (e.g. species, genera); plant damage (e.g. damage types) |
| **Link interaction strengths** | Frequencies of species-to-species encounters | Frequencies of plant damage variables (e.g., functional feeding groups, damage types, feeding event occurrences) |
| **Type of data used** | Presence/absence of plant host–arthropod and pathogen herbivore species | Presence/absence of plant host–arthropod and pathogen herbivore damage |
| **Establishing herbivory** | Determined by live insects caught in the act, laboratory rearing on hosts, or agricultural and entomological sources | Determined by observing arthropod and pathogen damage on plant organs |
| **Establishing generalization–specialization** | Determined on plant–host breadth of herbivore species | Determined by plant-host breadth of functional variables such as damage types |
| **Feature being examined** | Evolution of species-to-species interactions | Evolution of feeding strategies and herbivory patterns |
| **Applicability** | Modern record only | Modern and fossil records |
| **Dimensional domain** | Modern spatial dimension | Modern and fossil spatial and temporal dimensions |
| **Data limitation or advantages** | Primary data abundant (modern record); sampling issues | Primary data limited (fossil record); sampling issues |

Functional bipartite network-based analyses of plant and insect associations have been appearing in paleoecological contexts (e.g., Swain et al., 2021; Currano et al., 2021) and new work on applying them to modern assemblages is underway (Azevedo-Schmidt et al., in prep). One such recent approach is to use the most finely resolved metric of arthropod and pathogen damage, feeding event occurrences, that define an herbivore community on an individual plant species within a fossil plant assemblage (Xiao et al., 2022a–2022c, in prep). The characteristics of these functional networks (Table 1) offer a new, broadly applicable approach toward understanding the ecology of interspecific interactions. Even though myriad opportunities exist for using functional bipartite networks in both the fossil and the modern record, some concerns regarding the robustness of the network metrics and their applicability remain.

A major concern with various bipartite networks (including modern taxonomic ones) is that many of the patterns inferred from them may simply be artifacts of the sampling regime, and in particular, they may be a consequence of incomplete sampling (Hegland et al. 2010, Fründ et al., 2016), an issue that is also of concern for food webs (Paine, 1988; Wood et al. 2015). These problems are especially alarming when we are dealing with paleoecological assemblages (Shaw et al., 2021), where inherent bias due to differences in preservation, sample size, and collection methods present challenges to inference.

A host of methods exist that aim to statistically standardize comparisons among fossil plant assemblages. Most frequently, resampling has been done at the lowest sample size. Recent work (Currano et al., 2021) used resampling at 300 leaves as a procedure to investigate the metrics of bipartite networks constructed from angiosperm-dominated assemblages from the Late Cretaceous (ca. 67 Ma) to the recent (although they suggest sampling at higher numbers is always beneficial). In such resampling-based analyses, ecological specialization and structuring were found to be affected by major geological and ecological events such as the end-Cretaceous crisis, and by shifts in environmental variables such as temperature (Currano et al., 2021). These results are critical for decoding/understanding future climate change and the response of plant–insect interactions to future shifts in climate and environmental perturbations. Comparisons of functional bipartite networks in Currano et al. (2021) were completed using 500 bootstrap replicates for each assemblage resampled to 300 leaves, standardizing across uneven sampling regimes. Despite this consistency, however, the question remains of how the subsampling regimen affects the network values of the original assemblage. Here, we tackle this question by systematic reanalysis of data from 63 fossil plant assemblages representing almost all continents, varied paleolatitudes, and non-analog ecosystems (dataset of Currano et al., 2021). We explore the effect of resampling at different leaf sampling intensities (above and below 300 leaves) to determine the robustness of network metrics, assess paleoecologically relevant network-level and network class-level properties, and compare it to richness-based metrics that are often used in these systems (see Currano et al., 2021). We find that functional bipartite network representations not only perform better than many richness-based metrics, but also, with limited data, network metrics capture certain aspects of the complex structure of plant–DT interactions that are otherwise impossible to ascertain from aggregate richness metrics.

Overall, these efforts provide a partial resolution of the concerns associated with bipartite networks applied to functional plant–DT networks. Once the sampling impediments have been resolved, functional bipartite networks can illuminate the ecology of interspecific plant–insect interactions, which are not available in traditional taxonomic bipartite networks.

## Data and Methods

### Data Collection

The compiled dataset analyzed here was developed for a previous meta-analysis study (Currano et al. 2021). In that study, fossil insect damage census data published before 2021 were collected using Web of Science and Google Scholar using the keywords “insect herbivory”, “plant–insect interactions”, and “fossil”. Censuses selected for analysis were younger than 70 million years (Late Cretaceous) and dominated by dicotyledonous angiosperms. The sites were included in the meta-analysis only if they were collected in an unbiased manner; that is, only if all identifiable leaves were scored for herbivory and amounted to at least 300 dicot leaves. Each fossil assemblage dataset consisted of information about individual fossil leaves (e.g., taxonomic information) in the assemblage and a list of the number of different DTs present on each leaf. The fossil sites chosen represented a wide range of temporal, geographical, and environmental settings. For associated metadata referring to location and environmental conditions for each assemblage, consult the original work (Currano et al., 2021).

### Resampling

The basic independent unit of data/measurement in our work and related previous works is a single leaf and the associated DTs present on that leaf. As sample sizes (number of leaf specimens) differ among assemblages, this difference in sampling intensity can lead to spurious results while doing cross-comparisons, especially in metrics that heavily depend on sampling and this is especially true for network metrics (Hegland et al. 2010, Fründ et al., 2016; Costa et al., 2016; Schachat et al., 2018; Henriksen et al., 2019; Currano et al., 2021). Resampling all the assemblages to a lower common denominator number of leaves (i.e., rarefaction) is the accepted way of dealing with such a condition for a variety of richness and diversity-based metrics (see Schachat et al., 2018; Currano et al., 2021), but a detailed analysis of the robustness of such resampling on specifically network metrics remains unexplored.

To test the efficacy of sampling intensity on common network metrics, we resampled the data for each assemblage at the leaf level, using intervals relevant to the size of each assemblage (Table 2). For example, Chilga, an Oligocene site from Ethiopia, yielded 1063 leaves, and so we identified ‘resampling points’ every 50 leaves for the range 50 to 500 and at intervals of 100 leaves for the range 600 to 1000. The resampling stopped at 1000, even though the actual number of samples is 1063, for ease of comparison across assemblages at specified sample sizes. After finding these resampling points, resampling was repeated 5000 times. These intervals based on size of an assemblage were created to ensure reduced computational time as each run of network metric estimation was computationally expensive. The results do not reveal any new trends when done at finer resampling points for intervals of greater numbers of leaves but take a longer computational time.

**Table 2:** Sampling frequency at various ranges of sample sizes for the assemblages, and summary of number of assemblages within each sample size (specimens) grouping

| Sample size range | Sampling frequency | Number of assemblages with total leaves in this sample size range |
| --- | --- | --- |
| 0–500 | 50 | 18 |
| 500–1000 | 100 | 22 |
| 1000–2000 | 200 | 13 |
| 2000–4000 | 500 | 6 |
| 4000–8000 | 1000 | 2 |

### Metrics and comparisons

For each resampling of a given assemblage, we constructed a bipartite network using the occurrence data of DTs on individual leaves, such that plants and DTs are the two node classes in the network with edges or links connecting them. If one considers the adjacency matrix representation of this network, where rows are plant taxa and columns are DTs, we collated the number of DTs of a particular type on the leaves of a particular plant taxon in a single matrix element corresponding to the row and column represented by the plant and DT respectively. The reconstructed network was then used to calculate a set of metrics that have appeared in previous analyses of bipartite networks and may be ecologically meaningful metrics for plant–DT associations. These metrics have been used in previous works for different ecological systems (Bascompte and Jordano, 2013; Dormann et al., 2017), and in specific fossil plant–DT associations (Swain et al., 2021; Currano et al., 2021) to elucidate different aspects of structure of ecological networks, such as richness (number of species/DTs), prevalence of interactions (connectance), specialization (H2’, interaction evenness, partner diversity, togetherness), co-occurrence (C-score, niche overlap), resilience to random perturbations (resilience slope) and nestedness (traditional nestedness, NODF). These network metrics allow us to compare different aspects of the overall structure of different networks (Dormann et al., 2017), which otherwise are very difficult to compare directly (see Results section for more details on each metric in context).

We repeated this process of network metric estimation for each of the 5000 replicates for each possible resampling point of a given assemblage. To facilitate among-assemblage comparisons, we then normalized each metric using the value obtained from the network reconstructed from the complete data of an assemblage. In addition to an overall comparison, we divided the assemblages into five groups based on the number of samples (<500, 500–1000, 1000–2000, 2000–4000, and >4000 specimens; see Table 2 for a brief summary) as the number of specimens varied greatly among the assemblages of interest. We then examined how each network metric fared in the five groups at different sampling regimes using the normalized metric values obtained from the respective network reconstructed from the complete data of an assemblage.

## Results

From standard resampling procedures, it is expected that the resampled dataset would yield a lower richness than the full dataset, and this result is recovered (see Figure 1A–B). For both plants and DTs, a 75% recovery rate (i.e., the fraction of true value observed in a subsample) is observed for richness on average at the standard 300 sample size, and this continues to increase, as expected, with increased sampling. This relationship is the backbone of most of the comparisons using resampling in fossil plant–DT association studies (Currano et al., 2021). Given this result, one would expect that network metrics, which heavily depend upon the complex interactions among species, would perform even worse in resampling calculations as they depend highly upon the sampling and its accuracy. This is indeed true for a few widely used network metrics like functional complementarity (Figures 1C,D), which is a community-level measure of ecological niche complementarity and is measured as the total branch length of a (functional) dendrogram based on qualitative differences in DT presence among plants (Devoto et al., 2012). Functional complementarity was underestimated by about 50% at the 300 leaf sampling level, and this underestimation persisted even at higher sampling levels (Figure 1C,D). Given the sensitivity of dendrogram construction to the incorporation of new data, the result is not entirely surprising, especially for the elevated underestimation of rarer interactions. This does not mean that functional complementarity is not an interesting metric, but rather it is very sensitive to sample size and consequently it is not a practical metric to use in comparisons among assemblages.

**Figure 1:**
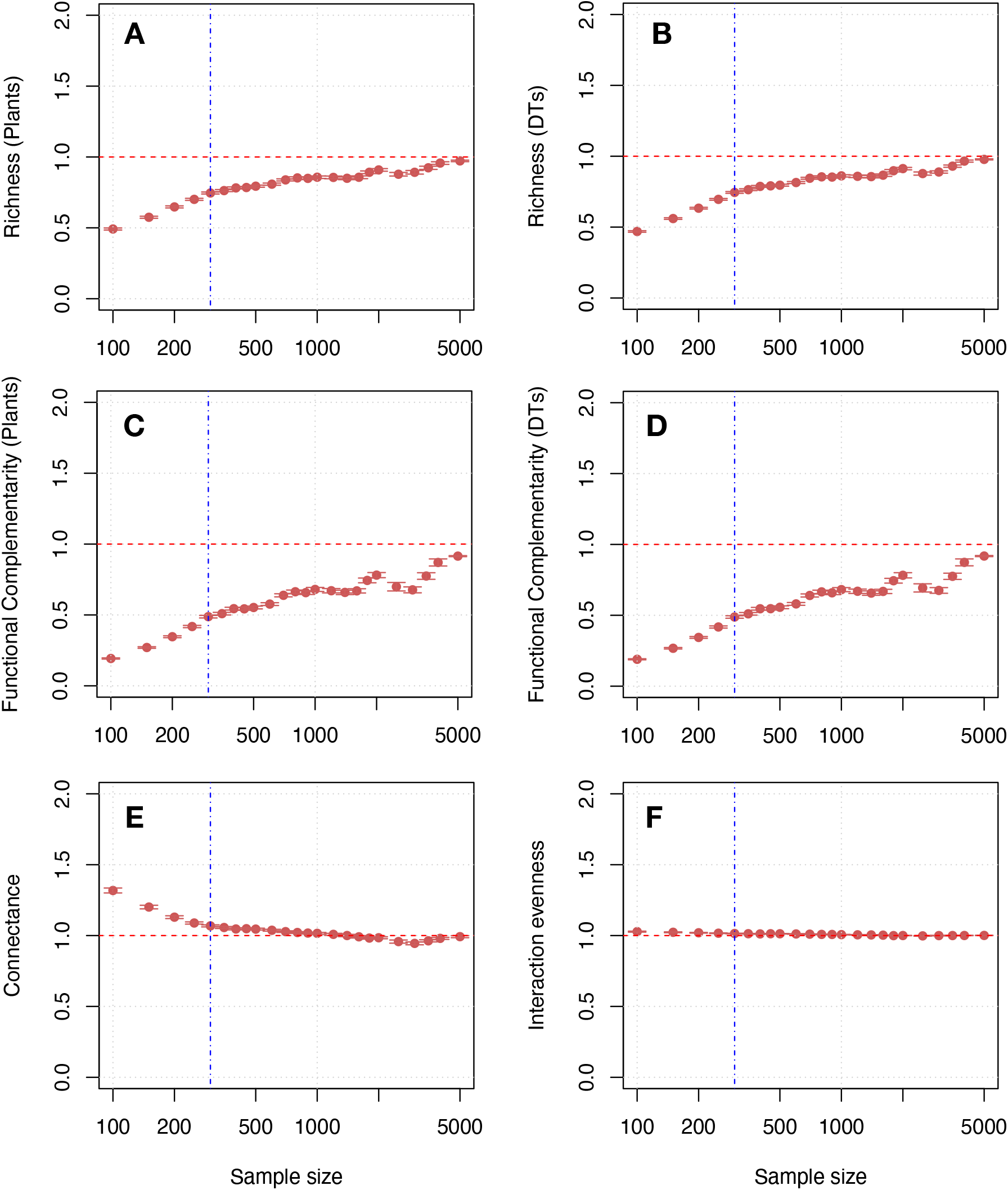
Normalized values of selected common network metrics averaged across all 63 assemblages (with 95% confidence intervals) at different sampling points. These metrics include the species richness of plants at (A); damage types (DTs) at (B); their respective functional complementarity at (C) and (D); connectance at (E); and interaction evenness at (F). The values for each assemblage, before being averaged, were normalized with respect to the metric value of the complete assemblage, thereby using all the available data for the assemblage, which would have a value of 1. The red dotted horizontal line denotes 1, and the blue vertical dotted line denotes the 300-sample size of leaves used in previous works (see Currano et al., 2021). Please note that the sample size axis is logarithmic.

Surprisingly, however, a large array of other network metrics do not depend as sensitively on sample size. For example, connectance, which measures the proportion of realized interactions to the total number of possible interactions, was overestimated approximately by a minimal 6% at 300 leaves (Figure 1E). Connectance was overestimated for most of the sampling regime because lower sample sizes routinely failed to include the rarer plant species and rarer DTs. Given such absences, connectance was artifactually inflated because there were fewer nodes in the resampled networks that could be linked together. As sample sizes increased (and assemblages became richer), the connectance shifted to being slightly underestimated (Figure 1E). This reversal from overestimation to underestimation occurred as ever larger levels of resampling included rarer plant species or DTs, but not necessarily the interactions with those nodes, thereby underestimating the realized number of interactions. Beyond 250 leaves, the connectance estimate remained, on average, within 10% of the value estimated with full data (VFD) for each assemblage (Figure 1E). This result is much better than the widely used richness metrics (i.e., the number of given taxa and related statistics) for plants and DTs, which hovered around 70% of the VFD estimate (Figure 1A-B) over a large range of sampling efforts. These differences among metrics as a function of sampling effort suggest that some aspects of network structure, like connectance, can reliably be understood at much smaller sampling efforts than can other metrics, such as traditional richness measurements.

The picture becomes even more interesting when we consider metrics that involve the number of interactions associated with each link. Such metrics require weighted edges/links in bipartite networks, rather than just presences or absences that involve unweighted edges/links in bipartite networks. Considering the weighted version of connectance, we found a 10% overestimation of connectance at 300 leaves (Figure S1), a percentage that is acceptable considering that the incorporation of abundance data can be very noisy. Another such example is interaction evenness (Tylianakis et al., 2007; Blüthgen et al., 2008), which characterizes how balanced the frequency of interactions is across the species and DTs in a given assemblage. (This metric can be thought of as akin to Shannon evenness.) The value estimate for this metric was within 2% of the value obtained from the VFD. Although interaction evenness is a metric that performs extremely well, one must be wary that presence–absence data usually is more robust than frequency data.

Indices related to network DT–host-plant (network) specialization showed similar promise (Figure 2). Niche Overlap, which measures the mean similarity of association patterns among DTs or plants of a given node class level (Figure 2 A,B), had an underestimation of about 6% for DTs and 10% for plants at the standard 300 leaf sample level. This underestimation is attributable to the incorporation of unknown associations that can increase the overlap between any two species/DTs. Togetherness, which is the mean number of co-occupancies across all species/DT combinations at a given node level, was under 5% error (at 300 leaves) for both DTs and plants, except for the special cases of highly diverse assemblages (discussed later; Figure 2 C,D). Togetherness for plants was slightly underestimated, as the full complement of interactions with rarer DTs usually are not incorporated, and the opposite pattern occurs in the case of DTs, where togetherness was slightly overestimated (Figure 2C,D). Partner diversity registers a 20–25% underestimation of the VFD at the 300 leaf sample level, which is appreciable, but still an improvement over the richness metrics (where the margin of error is much higher). Given the strict dependence of sample size on richness and abundance, these results are not surprising.

**Figure 2:**
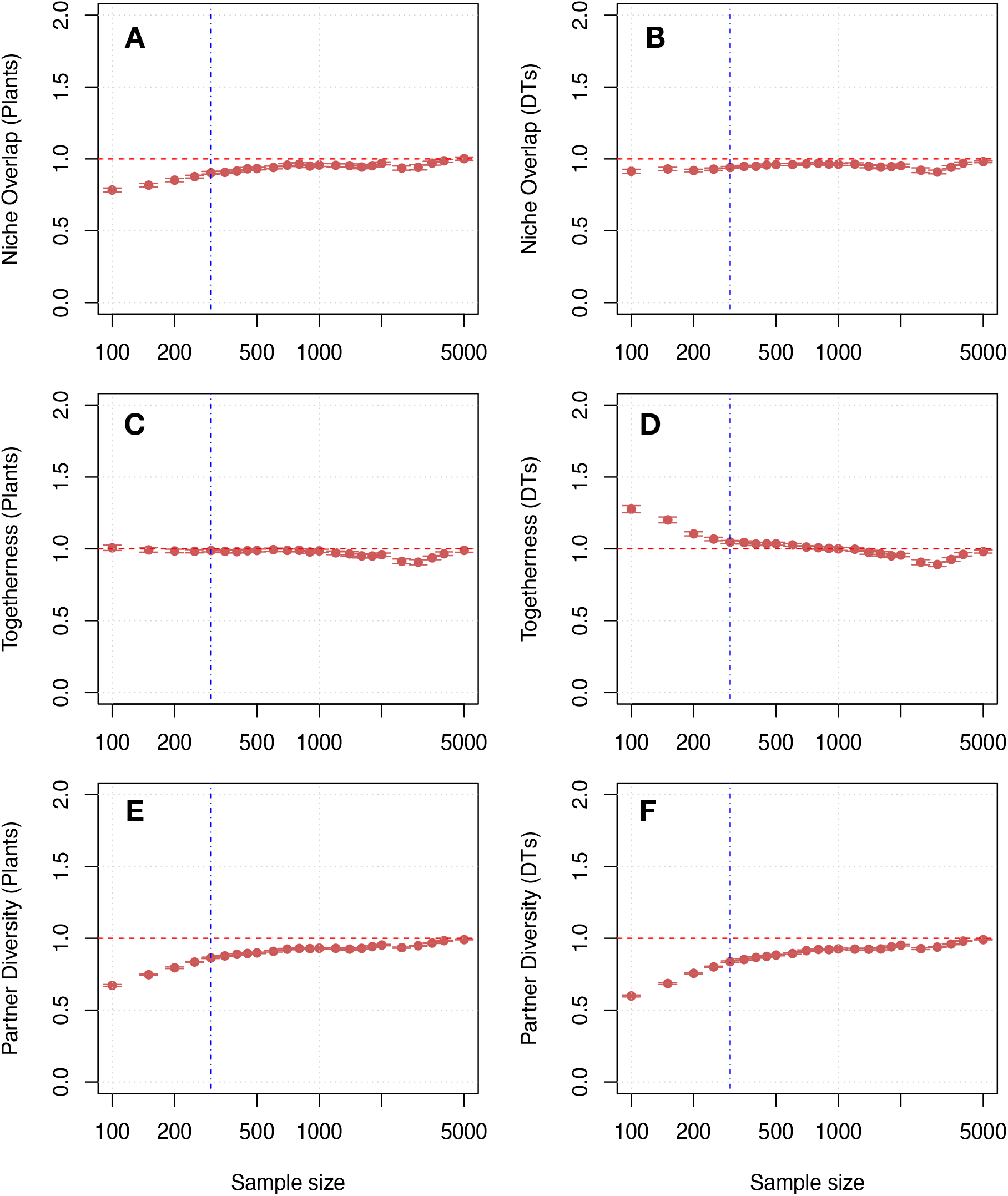
Normalized values of various specialization metrics averaged across all the assemblages (with 95% confidence intervals) at different sampling regimes. These metrics include Niche Overlap for plants at (A) and DTs in (B); Togetherness for plants in (C) and DTs in (D); and Partner Diversity for plants in (E) and DTs in (F). The values for each assemblage, before being averaged, were normalized with respect to the metric value of the complete assemblage, such that all the available data were used for the assemblage, which would have a value of 1. The red dotted horizontal line denotes 1, and the blue vertical dotted line denotes the 300-sample leaf size used in previous work (Currano et al., 2021). Please note that the sample size axis is logarithmic.

The C-Score, also called the Checkerboard score (Stone and Roberts, 1992), measures the mean number of checkerboard combinations across all nodes and indicates co-occurring associations. This metric was overestimated in both plants (by about 15% at 300 leaves) and DTs (by about 20% at 300 leaves; Figure 3 A,B), attributable to the incorporation of samples of rarer plants and DTs that can add an association previously considered as absent. Such an association is exclusive, or checkerboard-like to another association, and therefore reduces the C-Score.

**Figure 3:**
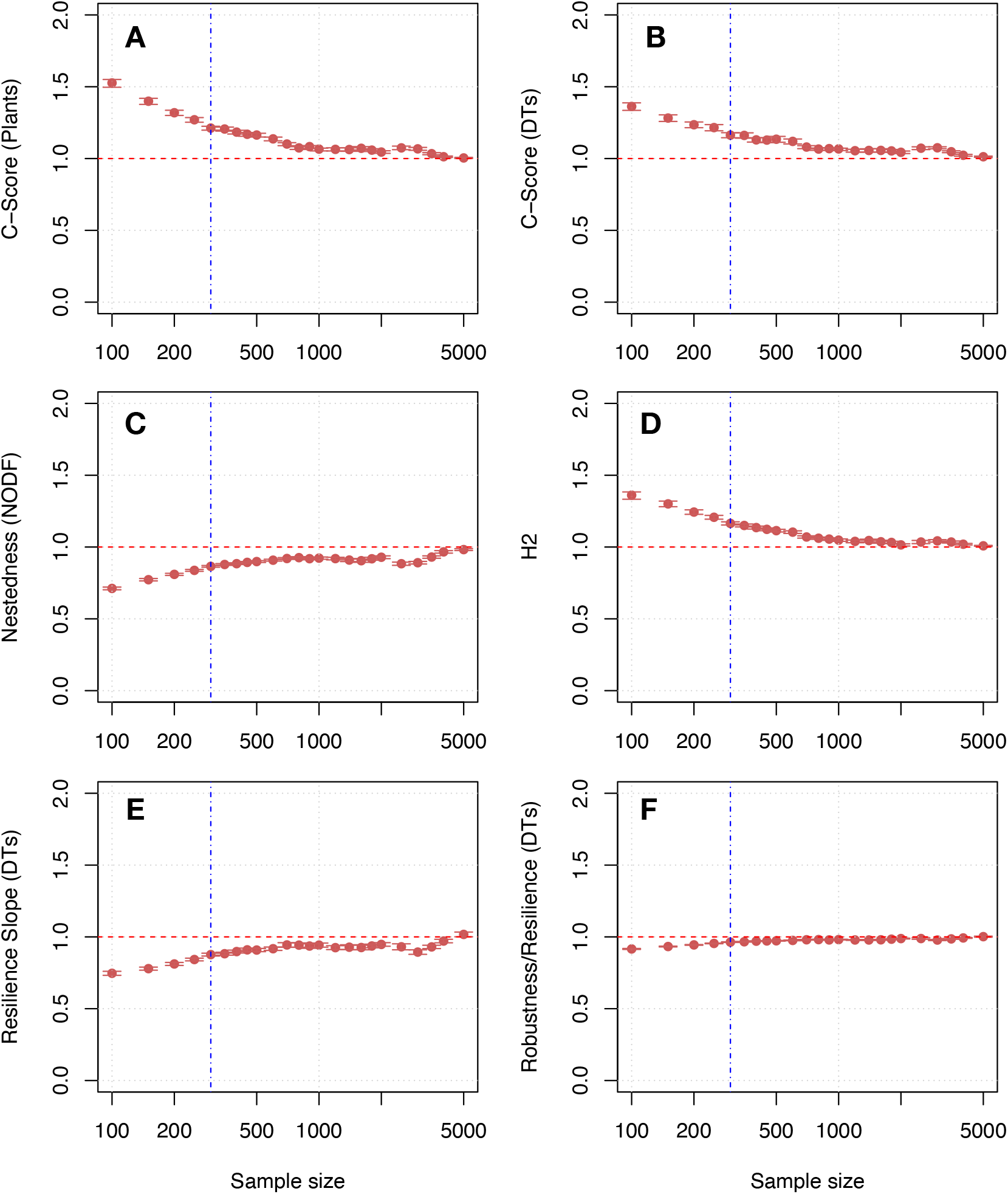
Normalized values of various network metrics averaged across all the assemblages (with 95% confidence intervals) at different sampling regimes. These metrics include the C-Score for plants at (A) and DTs at (B); Nestedness, or the Nestedness metric based on overlap and decreasing fill (NODF) at (C); H2, specialization at the level of the entire network, at (D); Resilience Slope (for DTs) at (E); and Robustness/Resilience (for DTs) at (F). The values for each assemblage, before being averaged, were normalized with respect to the metric value of the complete assemblage, using all the available data for the assemblage, which would have a value of 1. The red dotted horizontal line denotes 1, and the blue vertical dotted line denotes the 300 sample leaf level used in previous works (Currano et al., 2021). Please note that the sample size axis is logarithmic.

Measurement of nestedness in an ecological system can be accomplished in several ways (see Strona et al., 2014 for a brief overview). The most commonly used is the traditional metric of nestedness which is based on the ‘temperature’ of the occurrence matrix (BINMATNEST; Rodríguez-Gironés and Santamaría, 2006), and NODF, an acronym for nestedness metric based on overlap and decreasing fill (Almeida-Neto et al., 2008; Almeida-Neto and Ulrich, 2011). Estimations were calculated for both, and it was found that NODF performed much better than the traditional temperature-based metric (Figure S2); NODF underestimated the nestedness VFD by less than 15% at the 300 leaf sample level, whereas the traditional metric overestimated it by 50%. This makes NODF a better tool to measure nestedness in plant-host–DT bipartite networks, although it is generally less sensitive to randomness and data structure (Almeida-Neto et al., 2008). By contrast, the weighted versions of the nestedness metrics performed better using the traditional BINMATNEST methods than did NODF (see Figure S2).

H2’ is an often-used metric to assess how different are the interactions in an assemblage as compared to a random assortment; the assessment is based on the abundance values of both the plants and the DTs (Blüthgen et al., 2006). This metric was overestimated by 16% at the 300 leaf sample level as compared to the VFD. Given that H2’ uses abundance data for its estimation and not just presence-absence values, its performance is noteworthy but not surprising as in previous works. It has been shown to be robust to estimation based on partial data (Blüthgen et al., 2006).

In ecological bipartite networks, resilience slope, also termed the extinction slope, works on a repeated random sequence of plant node removals, and in our case calculates the number of secondary removals, such that if a DT is present in two plant species and both plants are removed, then the DT is not attached to any plant at the plant node level and is ‘secondarily’ removed, resulting in recalculation of its slope (Memmott et al., 2004). In network parlance, resilience/robustness refers to the area under this plot of secondary removal (Memmott et al., 2004), which in modern ecological texts is referred to as secondary extinctions, representing another measure of how tightly connected a network is. (See Zitnik et al., 2019 and Klein et al., 2021 for a broader network context for resilience). Here, we only consider plant removals as they provide the food source and can cause extirpation of DTs. The converse, pertaining to the impacts that the removal of functional DTs has on plant species, has an unclear meaning ecologically, and therefore is not used in this study. Resilience slope was underestimated by 13% and the ratio resilience/robustness was underestimated by less than 4% (at 300 leaves), making the latter a good metric for assessing the hardiness of assemblage structure to random secondary node removals (Figure 3E,F).

These network metrics, on average, addressed our ecological measurement needs and performed sufficiently well in the face of subsampling issues when averaged across all assemblages and their sample sizes. This can be seen from the fact that beyond 4000 leaves, the graph data trend lines became uneven for most of the metrics. This is because with only 3 assemblages larger than 4000 specimens there were too few assemblages for assessing metric performance in a comparative setting. Without a doubt, a range of sample sizes exists across assemblages, and hence the effects of subsampling would differ across sample sizes. To further investigate this, we plotted the variation of these network metrics relative to VFD for different sample size groups (see Table 2; Figure 4).

**Figure 4:**
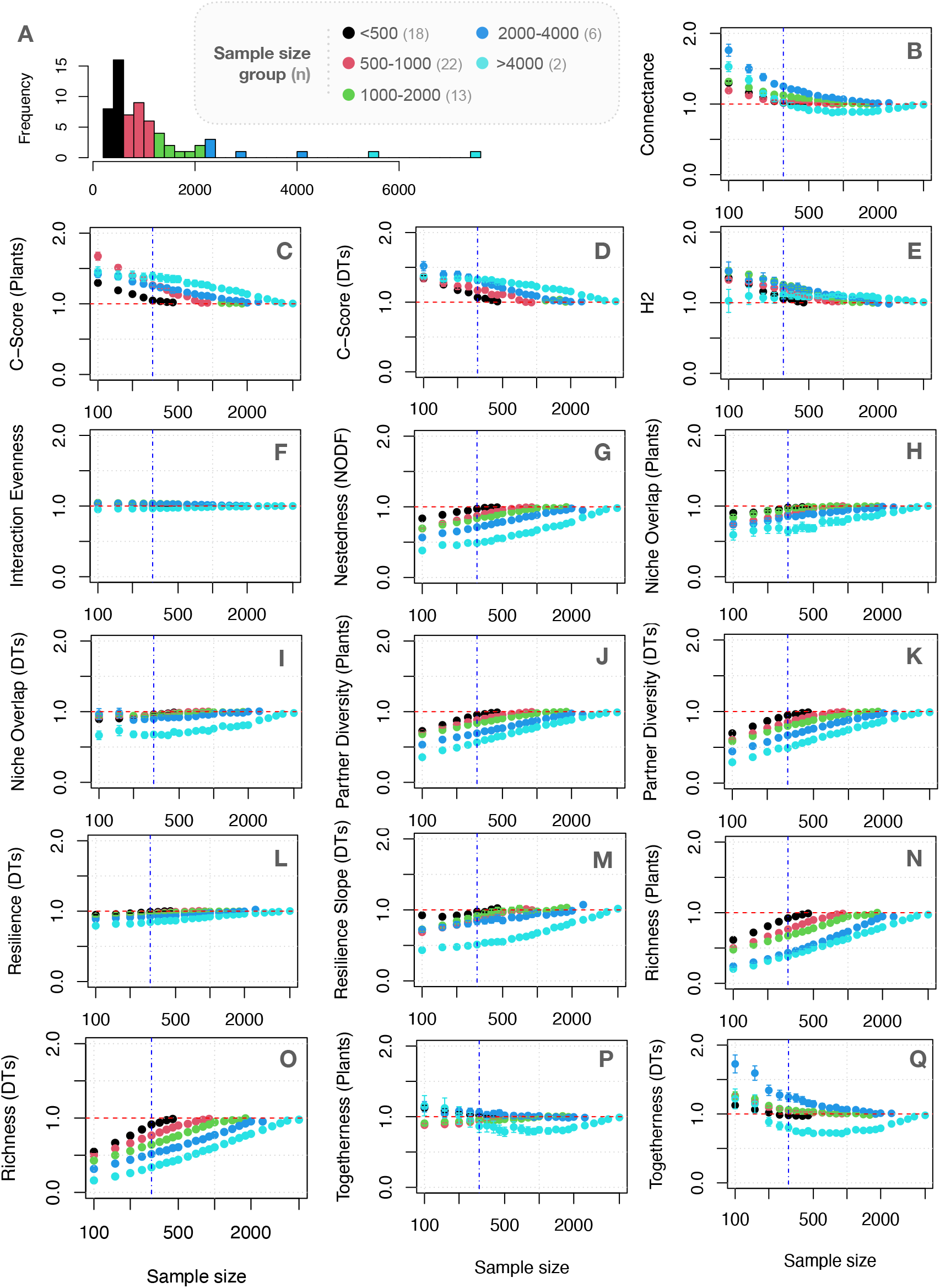
Normalized values of various network metrics averaged across all assemblages (with 95% confidence intervals) at different sampling regimes and sample size groups. The number of assemblages is indicated in each group. The values for each assemblage, before being averaged, were normalized with respect to the metric value of the complete assemblage, using all the available data for the assemblage, which would have a value of 1. The dotted red horizontal line denotes 1, and the blue vertical dotted line denotes the 300-sample leaf size used in previous studies (Currano et al., 2021). Please note that the sample size axis is logarithmic.

For ease of discussion, we will focus on the case of subsampling at 300 leaves for all metrics and the error from the VFD. Richness values for subsampled assemblages show a general expected trend of worse performance for larger sample size assemblages for a given subsampling size, with more than 50% underestimation for groups with 2000-4000 and >4000 leaves (Figure 4N,O). In contrast, network metrics that have performed well for the average case, continued to perform better across the subsampling range (Figure 4), with all of the metrics showing less than 50% under- or overestimation, and considerably better performance than richness. C-score, nestedness and partner diversity showed relatively worse behavior among the network metrics, but still performed better than richness (Figure 4). Metrics like connectance, interaction evenness, togetherness and resilience had less than 25% error from the VFD. This shows that selected network metrics reliably capture aspects of the interaction structure, even from partial data (subsampling).

## Discussion

Using functional bipartite network approaches, in contrast to taxonomic bipartite networks (Table 1), we sought to examine the robustness of network metrics to sampling intensity, and thereby assess the general usefulness of functional bipartite network approaches for the analysis of plant–insect associational data. Here, we show that many network metrics are reliable at estimating various interactions between plant and insect herbivores, using damage types (DTs; Labandeira et al., 2007) within paleontological datasets. Utilizing the functional network approach allows us to depart from taxonomic classifications, focusing on conserved functional traits/groupings, which are comparable across large swaths of geological time as well as geographical space.

Traditionally, studies of plant–DT associations have focused on quantifying the frequency and richness of DTs and Functional Feeding Groups (FFGs) from plant assemblages (Currano et al., 2021). Such determinations, while important, do not capture the intricate aspects of the interconnectedness and complexity of the feeding associations or aid understanding of potential drivers of change in a plant host–herbivore assemblage. Employing a functional bipartite network perspective affords a level of understanding about the complexity of such systems; however, whether the magnitudes of network metrics are factual representations of the systems or are consequences of sampling bias has been a major concern. Earlier, foundational works focusing on extant taxonomic bipartite ecological networks have explored the effects of undersampling on the robustness of network structure and associated metrics (Fründ et al., 2016; Vizentin-Bugoni et al., 2016). These studies have shown that taxonomic network metrics are differentially sensitive to undersampling and subsampling from both a theoretical perspective (Fründ et al., 2016) and with field data (Vizentin-Bugoni et al., 2016). The results reported here show the effectiveness and reliability of network metrics to capture aspects of the interaction structure of fossil plant-interaction networks in specific, and ecological networks in general (Figures 1–4). Through functional network-based methods, we find that even partial data about assemblages afford estimates that reasonably approximate the metrics of the overall assemblage. This makes the complex systems approach – where we look at the system as a whole without aggregating or averaging out intricate interactions – not only useful but also necessary.

Among the network metrics, connectance, interaction evenness, togetherness, and resilience performed the best across sampling regimes. Other metrics, such as H2’, NODF, C-Score, and Niche Overlap, performed better than richness, irrespective of the original sample size and subsampling range (Figures 1–4). Thus, our analyses identify a set of network metrics most likely to be robust to sampling effort. However, we caution that the precise nature of sampling effort in particular systems may deviate from the rarefaction methods we adopted here. Preservation of fossil leaves is a multiscale process that depends on a variety of environmental inputs such as climate, the site of the preserved plant assemblage, the local environment (during burial), vicissitudes of leaf damage, the chemical milieu of the plant specimens, and other factors (Greenwood and Donovan, 1991; Behrensmeyer et al., 2000; DiMichele and Gastaldo, 2008). Moreover, the observation of DTs is dependent on a fossil plant assemblage that has achieved or exceeded a threshold in which the plant specimens are sufficiently well preserved, abundant, diverse, and unbiased, and have voucher specimens deposited in an institution accessible to other researchers (e.g., Wilf, 2008; Gunkle and Wappler, 2015; Currano et al., 2021; Xiao et al., 2022a, 2022b; Maccracken et al., 2022). Developing better models that consider preservational bias, such as mode of preservation, will provide error margins for estimating interactions in a manner that is site- and taxon-specific, thereby allowing for quantification of site preservation quality. Expanding from simple rarefaction to incorporate such effects in subsampling routines may provide a better ecological signal, one that accounts for collection and various forms of preservational bias.

Bipartite networks for modern and fossil plant–insect associations often differ in sampling intensity, and modern Linnaean taxonomic networks use datasets derived from plants and insect herbivores instead of plants and DTs. Nevertheless, the plant–DT association data we considered is similar to modern data in two ways. First, interactions, as damage, are represented on leaves and other plant organs. Such types of damage are therefore proxy units for observation of insect herbivory, and more importantly, these DTs are similar across space and time, irrespective of species which lodged them. For bulk census data of modern floral assemblages, documentation of an interaction typically is independent of the existence of the two interacting species of a plant host and an interactive herbivore (Novotny et al., 2010; Oliveira et al., 2010). Rare exceptions occur when an insect is documented in the act of consuming its plant host in the field (Dyer et al., 2007; Salazar et al. 2018), or reared on their suspected hosts in the lab (Carvalho et al., 2014; Dyer, 1995; Dyer et al., 2007), options not available from fossil data, with rare exceptions such as scale insects on fossil plant organs (Xiao et al., 2021a). Second, leaves containing DT data from a fossil plant assemblage are both time-averaged (inherently representing multiple growing seasons) and space-aggregated, (representing elements of a regional flora and stable insect herbivore communities) (Kidwell and Flessa, 1995). Modern ecological studies focusing on similar patterns of plant–insect interactions with similar conditions of spatiotemporal averaging primarily focus on herbivory frequencies and diversities/richness (Adams et al., 2009; Smith and Nufio, 2004). These studies greatly differ from collection of foliage from a modern plant assemblage that also represents a local spatiotemporal sampling of habitats from a larger regional flora (Novotny et al., 2010) or insect feeding guilds sampled within a modern flora (Oliveira et al., 2020). Sampling of a fossil or modern plant assemblage, replete with time averaging and space aggregation, can be beneficial where there is a more comprehensive representation of plant taxa from adjacent local habitats within the regional flora (Burnham 1992; Olszewski, 1999), of which there are several examples in the fossil record (e.g., Xiao et al, 2021a–2022c). Paleobotanical leaf localities characterize spatiotemporal averaging with each site representing multiple growing seasons (e.g., Burnham et al. 1992). As these leaf samples are not from single growing events, they aid in our understanding of stable, non-random instances of plant–DT associations. This is important for characterizing stable communities through time, rather than herbivory at single plant assemblages. These similarities in the plant-host–DT data structure of fossil versus modern plant assemblages can provide continuity in the methods of assessing network metric robustness, although more work and data are needed for such comprehensive comparisons.

Due to the operational nature of DTs and the time-averaged and space-aggregated nature of DTs on plants, there is an important difference, at least preliminarily, between evaluating herbivory between fossil and modern assemblages. This difference involves how plant-host specificity of insects is determined from fossil plant assemblages versus how insect host-plant specificities are determined for modern insect host species. The DT plant-host specificity assignments of generalized, intermediate specificity, and specialized are based on the distribution of damage on species within a fossil plant assemblage. These assignments are analogous but not explicitly comparable, to the modern insect herbivore dietary categories of polyphagy, oligophagy, and monophagy, respectively (Xiao et al., 2021b, 2022a–2022c). Equivalencies or the lack thereof between analogous fossil and modern categories of host plant specificities would require a study of the kinds of DTs left on modern plants by the feeding activities of known insects with known host-plant dietary specificities.

Network metrics hold great promise in discerning the complexity of interactions between plants and insects that have dominated the terrestrial biodiversity landscape during the past 410 million years. Understanding this complexity through the lens of fossil plant–DT interactions can provide an exploration not only into the ecological structure of the herbivore communities of host-plant species, but importantly how plants and their insect herbivores have responded to major geological and evolutionary events such as mass extinctions at the Permian–Triassic and Cretaceous–Paleogene ecological crises (Labandeira et al., 2002, 2016; Carvalho et al., 2021), major transient shifts in the Earth’s climate such as the Paleocene–Eocene Thermal Maximum (Currano et al., 2008, 2010; Azevedo-Schmidt et al., 2019), and dramatic expansions of major plant groups such as seed plants during the Permian (Xu et al., 2018; Maccracken and Labandeira, 2020), angiosperm diversification in the mid-Cretaceous (Xiao et al., 2022a–2022c), and possibly the spread of grasses during the later Cenozoic. A promising development has been an extension of the FFG/DT system to modern floras in a variety of ecological studies (e.g., Adams et al. 2009; Smith and Nufio, 2007; Meineke et al., 2019). A current exploration is elucidation of functional herbivory networks on individual plant host species using the most highly resolved metric available – feeding event occurrences – for establishing variation in the structure of herbivory among plant assemblages (Xiao et al., 2022a–2022c, in prep). These and other studies include recording how DT richness is correlated with the diversity of their culprit leaf-chewing insects (Carvalho et al., 2014), evaluating the role of altitude on Mongolian oak component communities of insect herbivores (Sohn et al., 2019), assessing herbivory intensity along latitudinal gradients (Adams et al., 2009), determining the patterns of insect herbivory in invasive plants (Bachelot and Kobe, 2013; Beaulieu et al., 2018), and documenting increasing herbivory during the last century using herbarium specimens (Meineke et al., 2018). We expect such studies will continue to extend in both directions: characterizing additional, appropriate fossil plant assemblages—as well as modern plant assemblages—to address fundamental ecological questions that involve plants, insects, and their associations.

## Acknowledgments

This is contribution 402 of the Evolution of Terrestrial Ecosystems consortium at the National Museum of Natural History, in Washington, D.C. A Swain is funded by a James S. McDonnell Foundation (JSMF) postdoctoral Fellowship. SA Maccracken is funded by NSF PRFB award #2010800. ED Currano is funded by NSF EAR 145031.

## Data and Code availability

All data and code needed to reproduce this work is available at https://github.com/anshuman21111/resampling-fossil-leaves

## Supplementary Material

**Figure S1:**
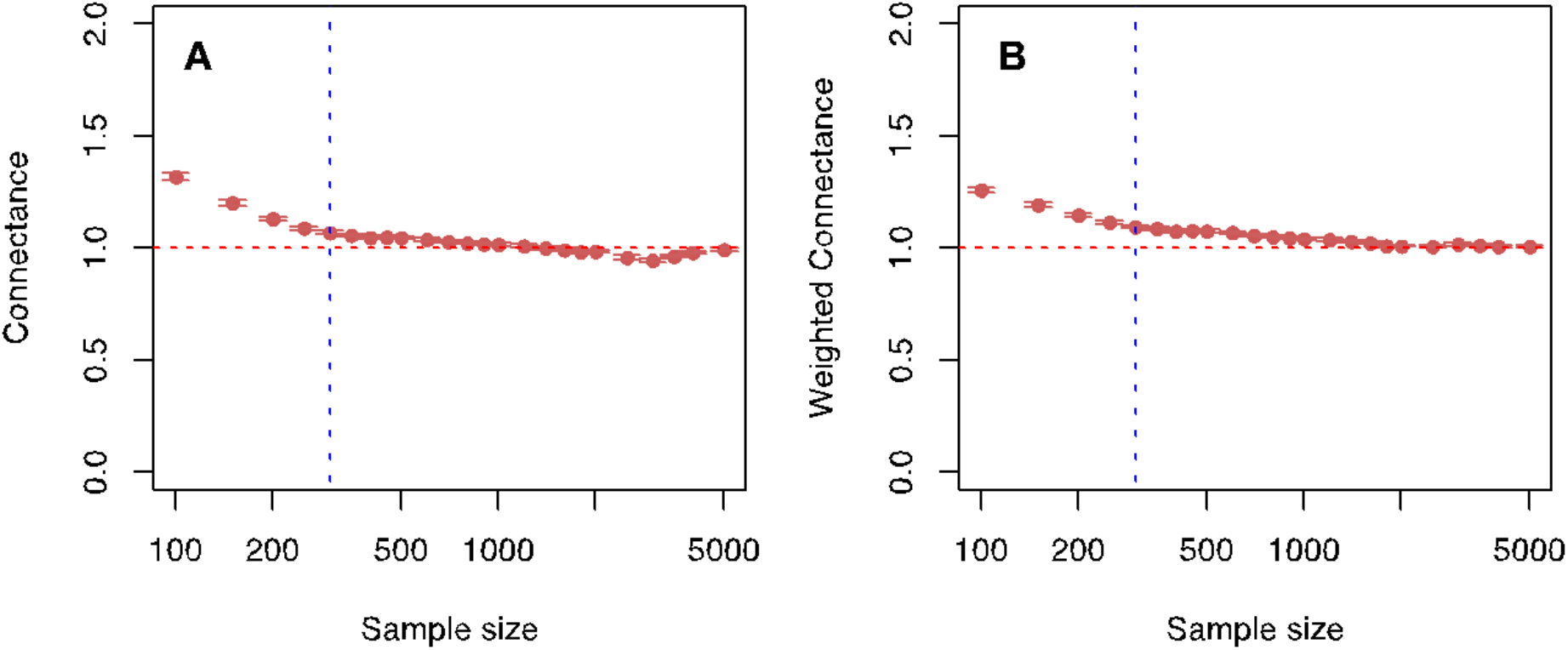
Normalized values of connectance (in A) and weighted connectance (in B) averaged across all the assemblages (with 95% confidence intervals) at different sampling regimes. The values for each assemblage, before being averaged, were normalized with respect to the metric value of the complete assemblage, using all the available data for the assemblage, which would have a value of 1. The red dotted horizontal line denotes 1, and the blue vertical dotted line denotes the 300 sample leaves used in previous studies (see Currano et al., 2021). Please note that the sample size axis is logarithmic.

**Figure S2:**
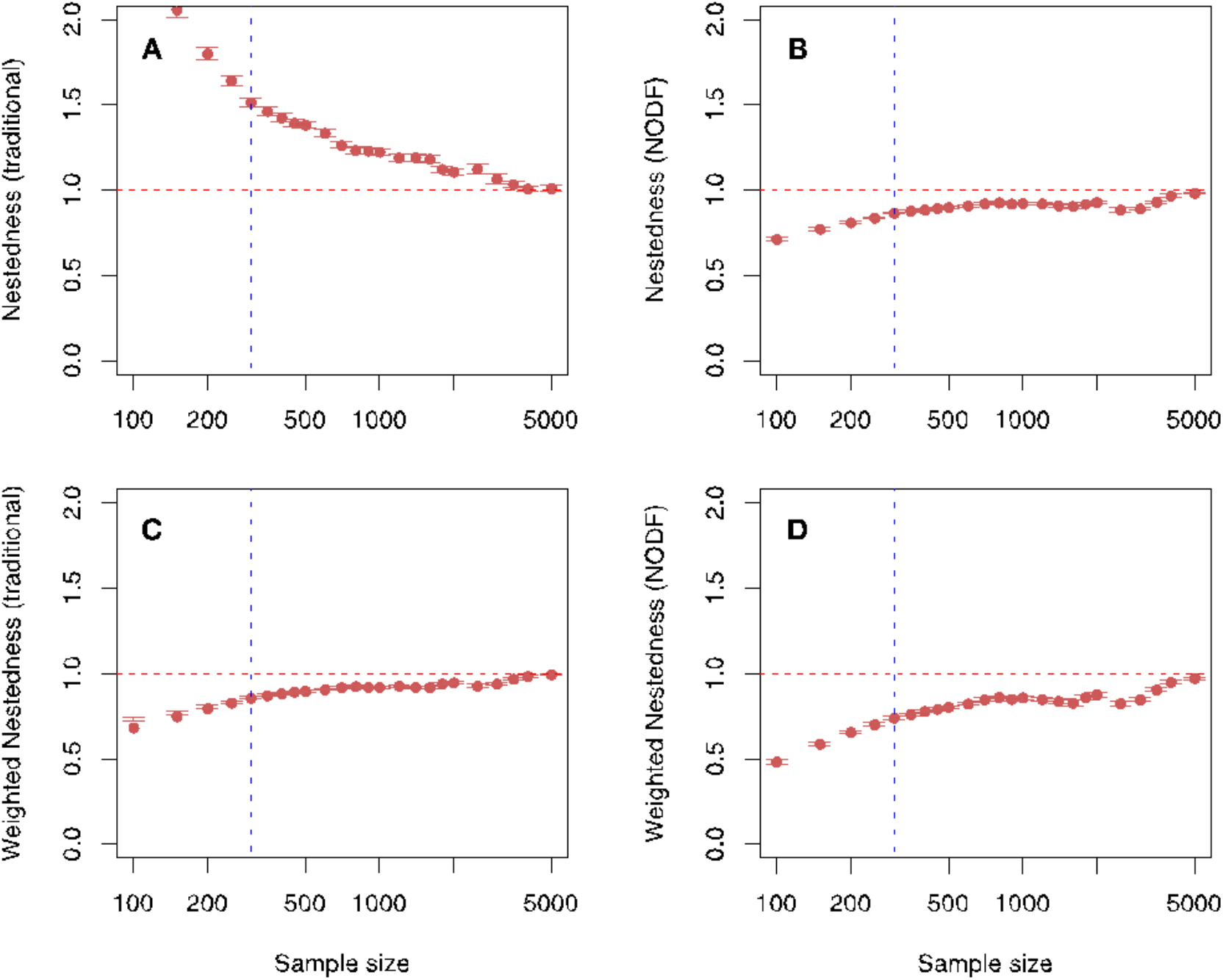
Normalized values of various Nestedness network metrics averaged across all the assemblages (with 95% confidence intervals) at different sampling regimes. These metrics include the Traditional Nestedness at (A); Nestedness (NODF or Nestedness metric based on overlap and decreasing fill) at (B), and their weighted counterparts in (C) and (D) respectively. The values for each assemblage, before being averaged, were normalized with respect to the metric value of the complete assemblage, using all the available data for the assemblage, which would have a value of 1. The red dotted horizontal line denotes 1, and the blue vertical dotted line denotes the 300 sample leaves used in previous work (Currano et al., 2021). Please note that the sample size axis is logarithmic.

